# Structure and function of the chromatin remodeler SMARCAD1 with its nucleosome substrate

**DOI:** 10.1101/2021.04.14.439859

**Authors:** Jonathan Markert, Keda Zhou, Karolin Luger

## Abstract

The ATP-dependent chromatin remodeler SMARCAD1 acts on nucleosomes during DNA repair and transcription, but despite its implication in disease, information on its structure and function is scarce. Chromatin remodelers use a variety of ways to engage nucleosomes, and outcomes of the ATP-dependent reactions vary widely. Here we show that SMARCAD1 transfers the entire histone octamer from one DNA segment to another in an ATP-dependent manner but is also capable of *de novo* nucleosome assembly from histone octamer, due to its ability to bind all histones simultaneously. We describe the cryoEM structure of SMARCAD1 in complex with a nucleosome and show that it engages its substrate unlike any other chromatin remodeler. Our combined data allow us to put forward a testable model for SMARCAD1 mechanism.

**One-Sentence Summary:** The single subunit chromatin remodeler SMARCAD1 engages nucleosomes in a unique manner and transfers the entire histone octamer.

---

DNA in eukaryotic cells is packaged by wrapping around octamers of the four core histones (dimers of H2A-H2B, and H3-H4) to form nucleosomes (*1*). These present a major barrier to DNA-dependent polymerases during transcription, replication, and repair. ATP-dependent chromatin remodelers slide, evict, or transfer (‘exchange’) histones to either space nucleosomes, to locally increase DNA accessibility, or to incorporate histone variants in place of major-type histones (*2*). The reactions catalyzed by the various remodelers, and their outcomes, depend on the type of remodeler and on the assays and substrates used. For example, SWR1 recognizes H2A containing nucleosomes, and in the presence of H2A.Z and ATP, removes H2A-H2B and replaces it with a histone variant dimer H2A.Z-H2B (histone exchange) (*3*). INO80, which is closely related to SWR1, slides nucleosomes along the DNA (*4*), while RSC slides nucleosomes but can also evict the histone octamer from DNA under certain conditions (*5*).

SMARCAD1 is a member of the INO80 family that is found throughout the genome and is implicated in DNA replication, transcription, and DNA damage repair (*6-14*). SMARCAD1 is of therapeutic interest due to its upregulation and/or mutation in pancreatic and breast cancer (*15, 16*). Human SMARCAD1 consists of two N-terminal CUE domains and a split catalytic domain (consisting of ATPase1 and ATPase2) which are connected by an extended loop that led to its classification into the INO80 family (*17, 18*). Virtually nothing is known about the structure and mechanism of human SMARCAD1, and only a few functional studies have been published with the *S. cerevisiae* and *S. pombe* homologues Fun30 and Fft3. Fft3 is found at open-reading frames of *S. pombe* genes and contributes to the eviction of histones from actively transcribed genes (*19*), while Fun30 and human SMARCAD1 appear to be primarily involved in histone eviction near sites of DNA damage to allow resection (*14, 20*). Fun30 has also been implicated in telomeric silencing in yeast (*6, 21, 22*). Fun30 has low nucleosome sliding activity, and instead evicts and exchanges histones between different DNA segments (*23*).

## SMARCAD1 prefers nucleosomes over free DNA for ATPase activation, and exchanges histone octamers

All chromatin remodelers studied to date require interactions with nucleosomes to activate their ATPase activity. To characterize the interactions and catalytic properties of SMARCAD1, we measured its affinity for a fluorescently labeled nucleosome reconstituted onto a 165 base pair (bp) DNA fragment containing the 147 bp 601 positioning sequence (7N11; numbers indicate the bp of extranucleosomal DNA), using a fluorescence polarization (FP) assay (fig. 1A, S1). Binding to an Alexa-488-labelled 147 bp DNA fragment was also determined. SMARCAD1 binds nucleosomes and DNA with similar affinities (K_D,Nuc_ = 160 ± 10 nM and K_D,DNA_= 80 ± 10 nM).

**Fig. 1:**
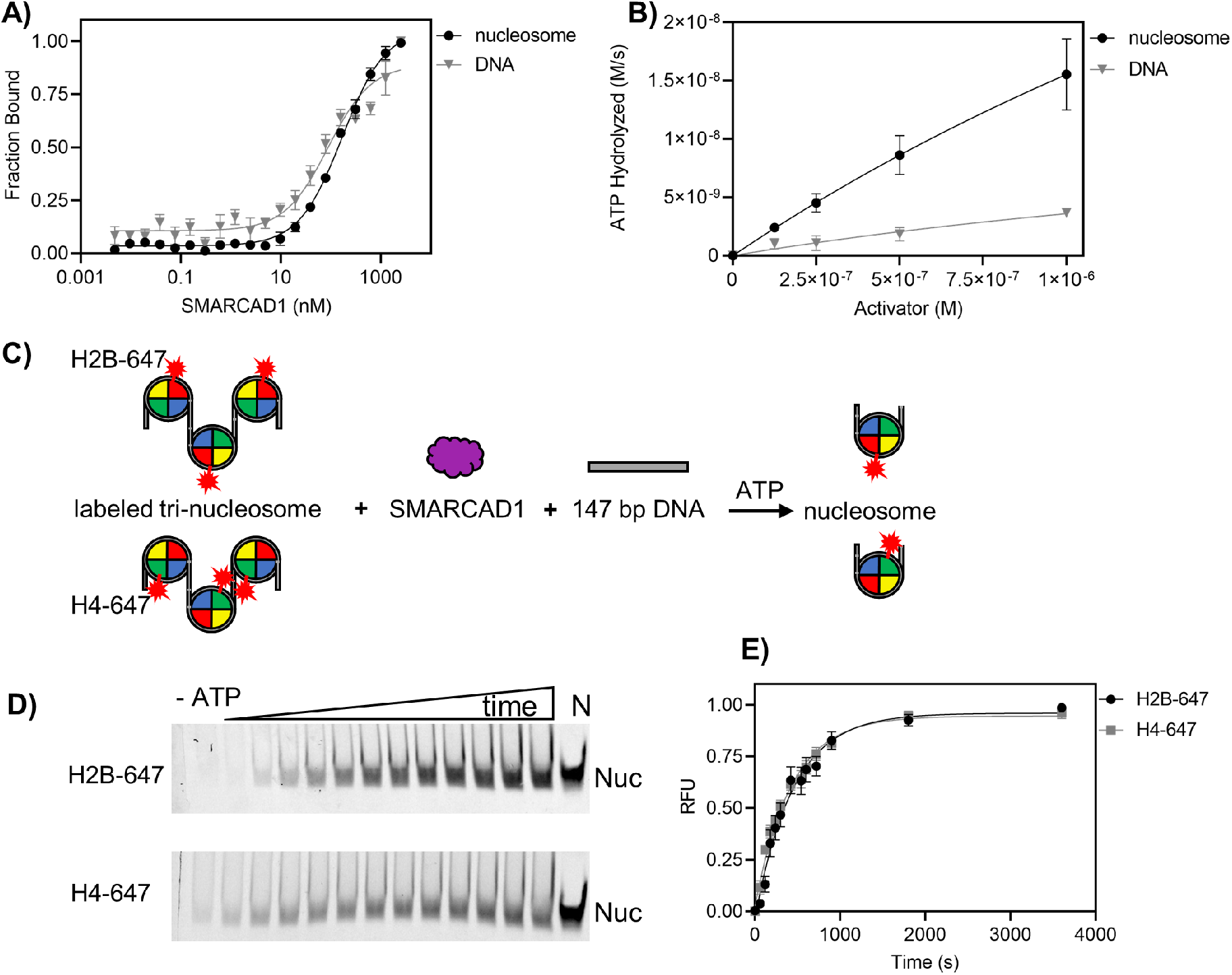
SMARCAD1 is an ATP-dependent chromatin remodeler with histone exchange activity. **A)** Fluorescence polarization assay of SMARCAD1 (0-2,500 nM) with 10 nM Alexa-488-labeled 7N11 nucleosome or 10 nM 488-labeled 147 bp DNA. K_D,nuc_ = 160 ± 10 nM; K_D, DNA_ = 80 ± 10 nM. K_D_ and standard errors (s.e.) were determined from three replicates. **B)** ATP-hydrolysis assay of SMARCAD1 (100 nM) with 7N11 nucleosome (0-1,000 nM) or 165 bp DNA (0-1,000 nM). S.e. (bars on graph) were determined from three replicates. **C)** Schematic of histone-exchange assay. SMARCAD1 (3 μM) was mixed with Atto-647 labeled tri-nucleosomes [(30N60N60N30] (75 nM tri-nucleosome / 225 nM mono-nucleosome) and 147 bp acceptor DNA (1.5 μM). Reactions were initiated with the addition of 1 mM ATP and quenched at indicated time-points (fig. S2) with EDTA followed by visualization on a 5% TBE-native gel. **D)** 5% TBE-native gel of histone-exchange assay (described in C) shows that in the absence of ATP (for 60 minute time-point), little H2B-647 or H4-647 is exchanged. When ATP is added and reaction is allowed to proceed 0-60 minutes, there is an increase in exchange, and product corresponds to nucleosome control. **E)** Quantification of D). SMARCAD1 exchanges both Atto-647 H4 (k_obs,H4_ = 2.2 ± 0.2 x 10-3 s-1) and Atto-647 H2B (k_obs,H2B_ = 2.3 ± 0.1 x 10-3 s-1) at the same rate. k_obs_ and s.e. were obtained from five replicates.

An ATP-hydrolysis assay that couples ATP-hydrolysis to NADH oxidation was used to compare the ability of nucleosomes and DNA to activate SMARCAD1 ATPase activity (*4*) (fig. 1B). This assay reveals that despite the similar affinity for DNA and nucleosomes, the latter are much more potent in activating SMARCAD1 catalytic activity.

*In vitro*, Fun30 promotes histone exchange rather than sliding (*23*), while Fft3 evicts histones in actively transcribed regions (*19*). To test whether SMARCAD1 exhibits histone exchange activity, we prepared tri-nucleosomes (fig. 1C) with either H2B or H4 carrying a fluorescent label. SMARCAD1 and 147 bp acceptor DNA were added, reactions were initiated by the addition of ATP, and quenched with EDTA after the indicated times. The rate of histone transfer from tri-nucleosomes onto 147 bp acceptor DNA was determined by quantifying the appearance of a fluorescently labeled mono-nucleosome (fig. 1D, E, S2). In the absence of ATP, no histones were transferred, while in presence of ATP, both H2B and H4 were transferred onto the acceptor DNA to form particles that migrate like the nucleosome control. Intriguingly, both H2B and H4 are transferred to form mono-nucleosomes at identical rates (fig. 1E), suggesting that the two histone dimers are transferred together.

## SMARCAD1 binds histones in absence of DNA and functions as a nucleosome assembly factor

Our data predicts that SMARCAD1 should bind all histones simultaneously in the absence of DNA, and this was tested using Fluorescence Resonance Energy Transfer (FRET). Histones H3-H4 and H2A-H2B do not interact under physiological conditions in absence of DNA. Refolded histone dimers (H3-H4* and H2A-H2B*, labeled* with Alexa-488 and Atto-647, respectively) were combined at low concentrations, and incubated with increasing amounts of SMARCAD1 (fig. 2A). The observed increase in FRET (K_D,histones_ = 2.1 ± 0.1 nM) reveals that SMARCAD1 brings together histones H2B and H4 in the absence of DNA (Fig. 2B). We also tested the ability of SMARCAD1 to bind H3-H4 or H2A-H2B on their own. H4 and H2B were both labeled with an Alexa-488 fluorophore, and SMARCAD1-histone binding was monitored in an FP assay. These assays revealed that SMARCAD1 binds H3-H4, and H2A-H2B independently (K_D,H3-H4_ = 1.2 ± 0.4 nM and K_D,H2A-H2B_ = 10 ± 2 nM) (fig. S3).

**Fig. 2:**
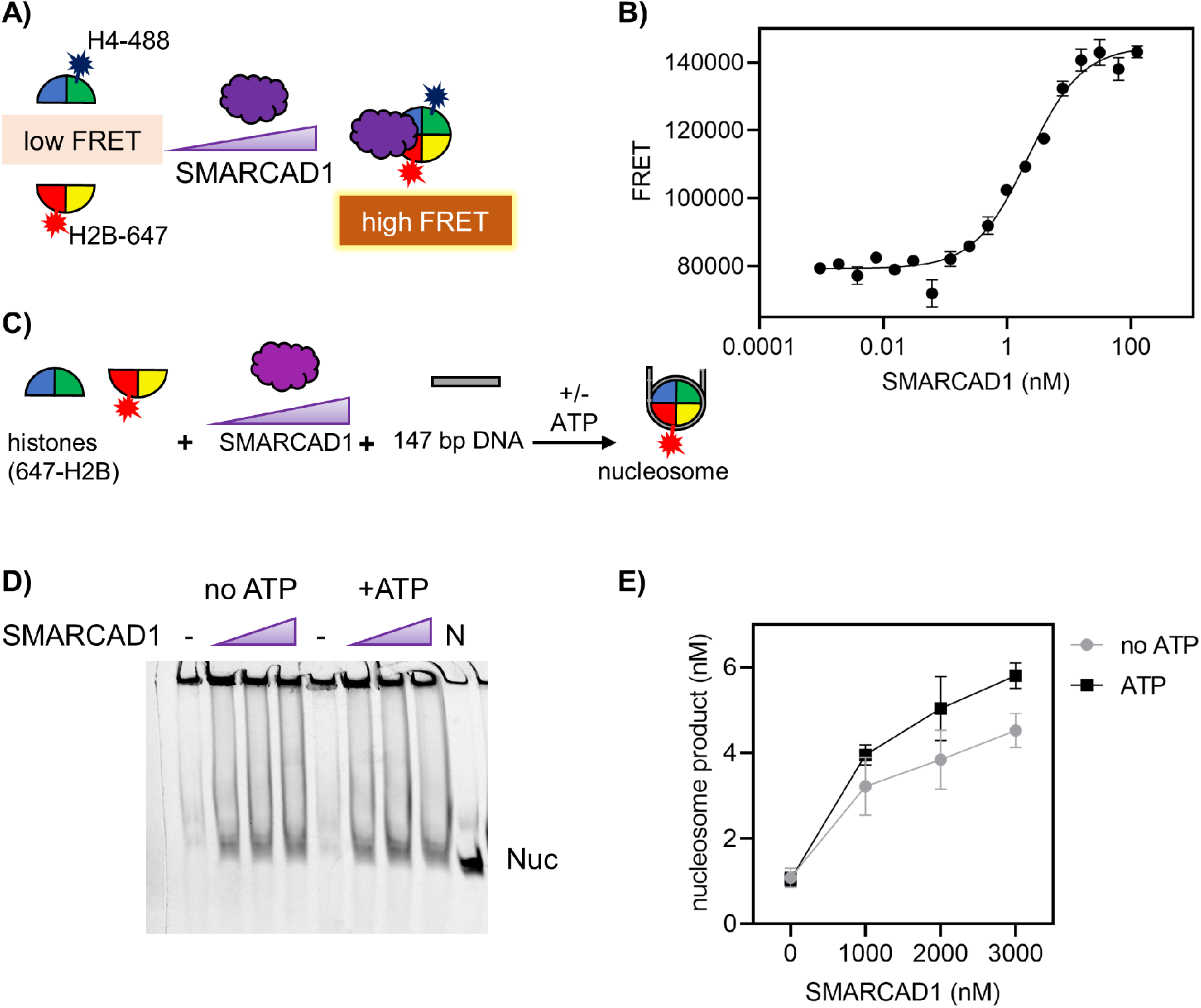
SMARCAD1 binds histones and assembles nucleosomes independently of ATP. **A)** Schematic of FRET-based histone binding assay. To monitor histone binding, H3-H4 (Alexa-488 H4) and H2A-H2B (Atto-647 H2B) were mixed at 10 nM. SMARCAD1 was titrated to histones (0-125 nM), and the increase in FRET signal is measured. **B)** SMARCAD1 binds histones in absence of DNA with high affinity (K_D,histones_ = 1.3 ± 0.4 nM). K_D_ and s.e. were obtained from three replicates. **C)** Schematic of nucleosome assembly reaction. H3-H4 and H2A-H2B (50 nM; 647-H2B) are mixed with SMARCAD1 (0-3 μM) for 15 minutes followed by addition of 147 bp DNA (50 nM) which was incubated for a further 15 minutes with and without 1 mM ATP. Reactions were quenched with pUC-19 plasmid DNA and then analyzed on a 5% Native-TBE gel, shown in **D)**. In the absence of SMARCAD1, no histones are assembled onto DNA; an increase in nucleosome assembly is observed upon addition of SMARCAD1. Nucleosome assembly does not require ATP. **E)** Quantification of gel from D); s.e. bars in graph are from three replicates.

Results shown in figures 1D, E indicate that SMARCAD1 transfers histone octamer from one DNA fragment to another. To test if SMARCAD1 promotes *de novo* nucleosome assembly (without first taking them off another DNA fragment), we pre-incubated SMARCAD1 with a mixture of H2A-H2B (Atto-647) and H3-H4, then added 147 bp DNA (fig. 2C). The appearance of fluorescent nucleosomes on a native gel was then quantified. In the absence of SMARCAD1, only minor amounts of nucleosomes were formed. As SMARCAD1 is added, the nucleosome band intensifies (fig. 2D, E). Unlike for the reaction shown in figure 1, where histones must first be removed from the DNA in an ATP-dependent manner, ATP is not required for the nucleosome assembly reaction that starts with free histone complexes.

## SMARCAD1 interacts with nucleosomes in a unique manner

We determined the cryoEM structure of SMARCAD1 bound to a nucleosome at an overall resolution of ∼6.5 Å (fig. S4). In our first screens, we were unable to observe SMARCAD1 density on a 147 bp nucleosome, and upon optimization ultimately used a nucleosome substrate with 70 bp extranucleosomal DNA extending from one side (0N70). To further stabilize the complex, we used GraFix with glutaraldehyde (*24*). In addition, a non-hydrolysable ATP analog (prepared by combining ADP with BeSO_4_ and NaF to form ADP-BeF_3_) (*25*) further stabilized the complex.

The density for the nucleosome was well-defined and could be visualized at ∼5 Å, and placing the human nucleosome (PDB 2CV5) was straightforward. We also observed density for ∼20 of the 70 bp extra-nucleosomal DNA which we built manually using Coot (*26*). The density for SMARCAD1 was at ∼10 Å and thus we were unable to confidently build a high-resolution model of SMARCAD1, as no structure of SMARCAD1 in isolation is available. We used Swiss-Model to predict a structure of SMARCAD1 using Iswi (PDB 5JXR) as a template (*27*). This model was then docked into the EM map using Chimera (*28*), and run through Flex-EM (*29*). The model was simulated in MDFF (using the NAMD engine and visualized in VMD) and further processed in ISOLDE to increase model to map fitting (*30-33*). Even though we used full length SMARCAD1 for grid preparation, we were able to see only density corresponding to the C-terminal catalytic domain encompassing ATPase 1 and 2 (fig. 3A).

**Fig. 3:**
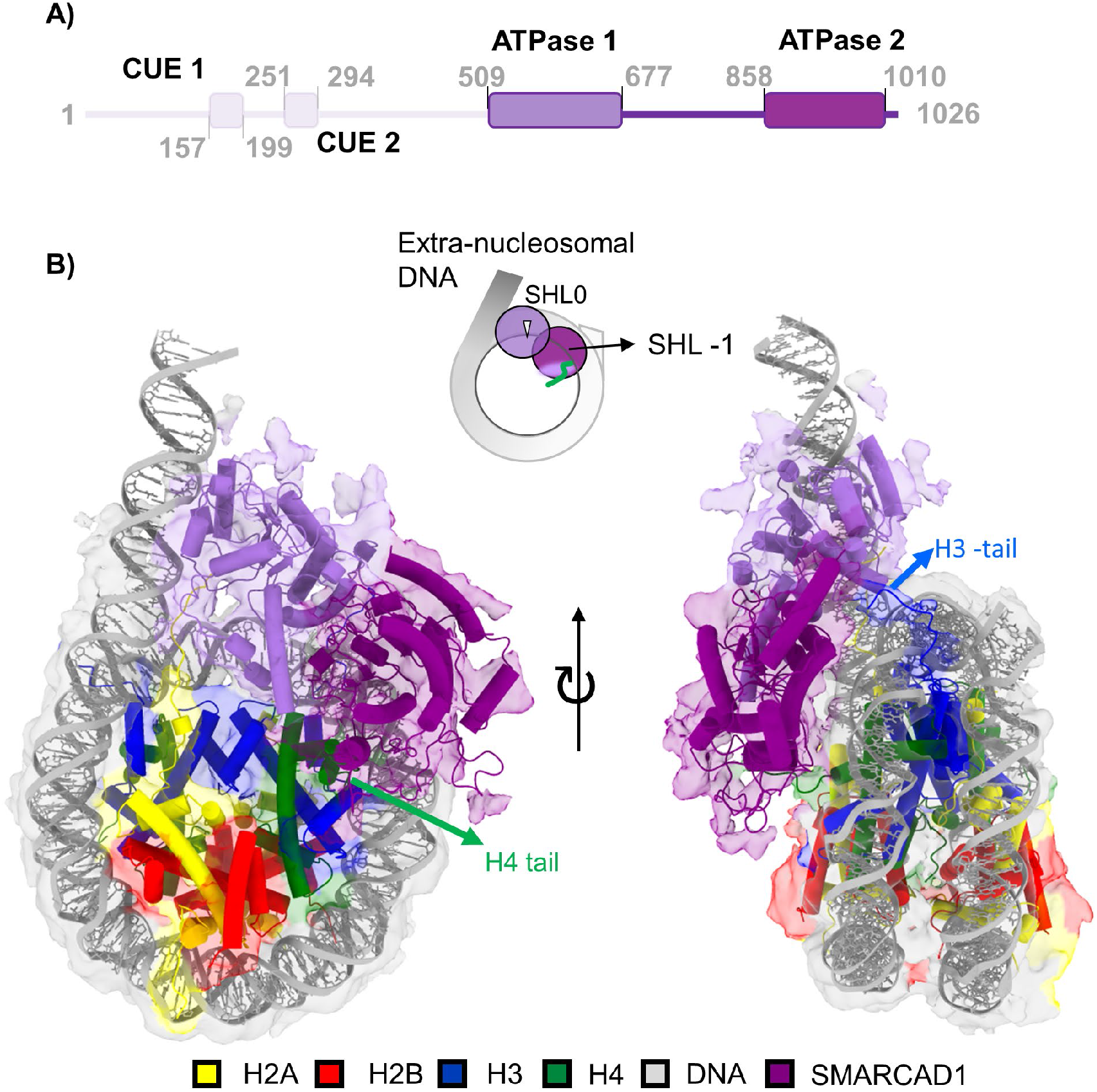
CryoEM structure of the SMARCAD1-nucleosome complex, revealing a unique binding mode. A) domain structure of SMARCAD1, with the domains that are visible in the density map shown in purple. **B)** SMARCAD1 makes interactions with extra-nucleosomal DNA, the nucleosomal dyad (SHL 0), and SHL -1, as well with the H4 N-terminal tail and the H3 N-terminal tail. Although full-length SMARCAD1 was used for cryoEM sample preparation, only the C-terminal catalytic domain (amino acid 509-1026) was visible in the map.

The low resolution of SMARCAD1 precludes interpretation of the molecular interactions, but we can draw conclusions on the overall placement of SMARCAD1 on the nucleosome (fig. 3B). The C-terminal catalytic domain of SMARCAD1 interacts with the nucleosome at several distinct contact points (DNA locations on the nucleosome are defined by their superhelix location, or SHL, where SHL 0 denotes the nucleosomal dyad (*1*)). SMARCAD1 sits on top of SHL 0 where it covers ∼10 bp of nucleosomal DNA. ATPase1 interacts with ∼10 bp of the extranucleosomal DNA and reaches towards SHL 0. ATPase2 continues to interact with the dyad DNA, but also maintains contacts with DNA near SHL -1 (movie S1). The remaining ∼500 amino acids of SMARCAD1 for which no density is observed, is predicted to be disordered. These are directed over SHL 0 towards the distal side of the nucleosome. The overall architecture of the nucleosome is unchanged compared to unbound nucleosome in the main class (fig. 3B).

We identified a second class of particles at ∼4.4 Å (map 2) (fig. S4). In this class, there is no discernable difference in how SMARCAD1 interacts with the nucleosome, nor are there any structural rearrangements in the histones, but nucleosomal DNA on the distal side of SMARCAD1 is peeled away from the histone core starting near SHL +4 (fig. S5).

## SMARCAD1 requires both H3 and H4 N-terminal tails for catalytic activity

The catalytic domain of SMARCAD1 appears to make direct contacts with the H4 and H3 N-terminal tails (fig. 3B, 4A, B, movie S1), although the molecular details of the interaction remain to be determined. In light of this finding and given the established importance of histone tails in regulating the activity of other chromatin remodelers (*2*), we tested their contribution to SMARCAD1 catalytic activity. SMARCAD1 binds nucleosomes reconstituted with tailless histones H3 or H4 (TLH3 and TLH4) with the same affinity as it binds nucleosomes with full length histones (fig. 4C). However, ATP hydrolysis is significantly reduced for TLH4 nucleosomes (fig. 4D), and the histone-exchange activity for TLH4 nucleosomes is abolished (fig. 4E, F, S6). In contrast, removal of the H3 tail does not affect ATP hydrolysis (fig. 4D). Nevertheless, the relative amount of histone exchange is significantly reduced in TLH3 compared to wild type (fig. 4E, F, S6). To ensure that the lack in exchange activity in TLH3 and TLH4 is not simply due to the inability of SMARCAD1 to interact with histones, we demonstrate that SMARCAD1 assembles TLH3 and TLH4 nucleosomes from histone complexes (de novo nucleosome assembly activity; fig. S7).

**Fig. 4.**
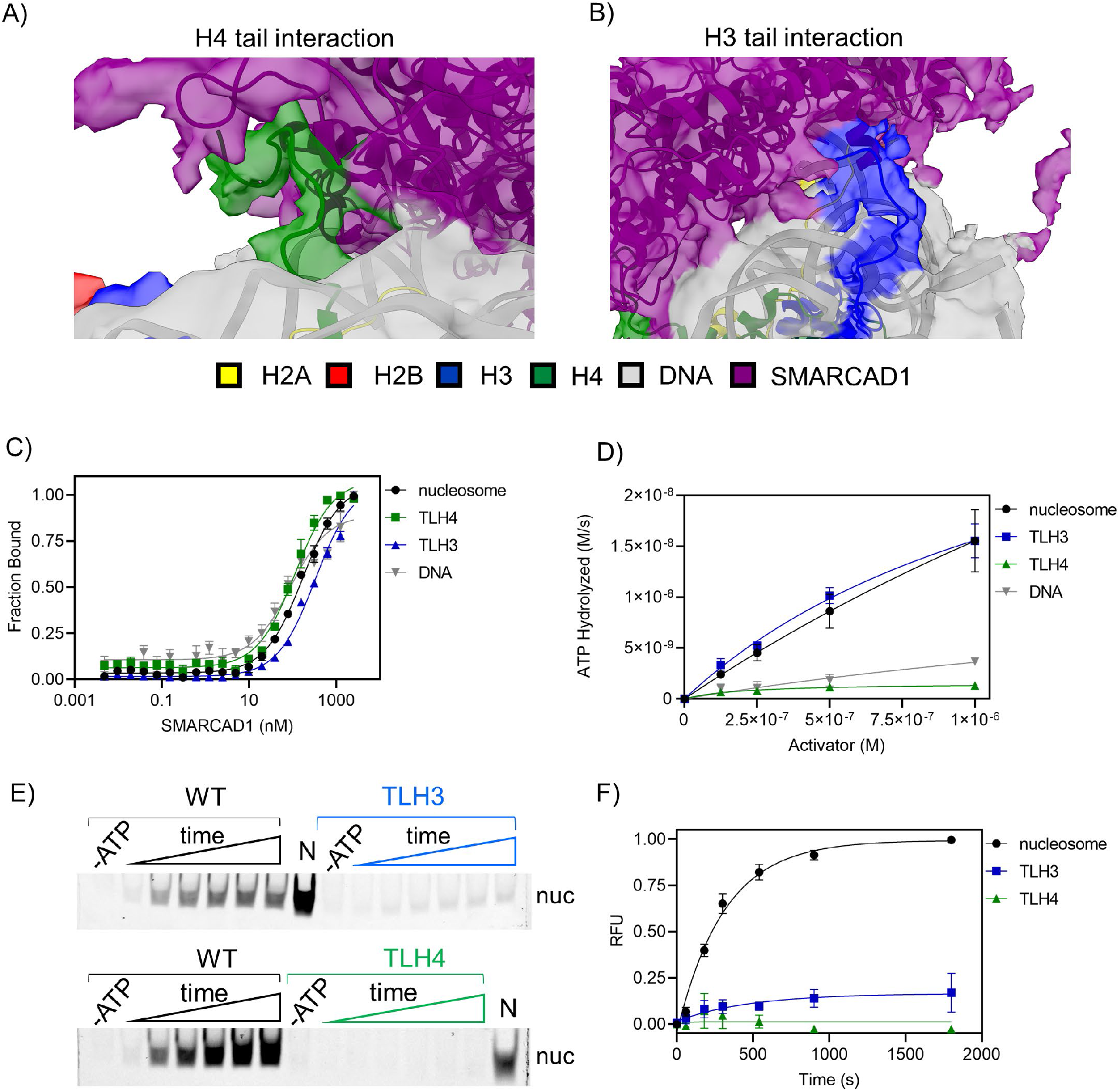
Histone H3 and H4 tails are important for SMARCAD1 activity. **A)** SMARCAD1 makes extensive contacts with the H4 tail. CryoEM map (transparent) with model reveals that SMARCAD1 makes several contacts with the H4 tail and **B)** with the H3 tail. **C)** SMARCAD1 binds nucleosomes with full length histones (WT), as well as nucleosomes with tail-deleted histones (TLH4, TLH3), and DNA. K_D,nucleosome_ = 160 ± 10 nM, K_D,TLH4_ = 110 ± 10 nM, K_D,TLH3_ = 330 ± 20 nM, K_D,DNA_ = 80 ± 10 nM. K_D_ and s.e. were obtained from three replicates, data for WT nucleosome and DNA are same as in figure 1A. **D)** SMARCAD1 is activated by WT nucleosome and TLH3 nucleosomes equally well, but TLH4 nucleosome and DNA are poor activators of ATP-hydrolysis. s.e. bars are from three replicates, data for WT nucleosome and DNA are same as in figure 1B and included for clarity here. **E)** histone exchange assay reveals that SMARCAD1 exchanges WT histones well, but it is unable to exchange TLH3 or TLH4 histones. Time points were 0, 60, 18, 300, 540, 900, 1800 seconds. **F)** Quantification of gel fromE) confirms SMARCAD1 poorly exchanges TLH3 or TLH4 histones compared to WT histones. s.e. bars in graph are from three replicates (6 for wild type nucleosomes).

### Discussion

ATP-dependent chromatin remodelers engage with their nucleosome substrate in a variety of ways, but, with the exception of INO80 and SWR1, the ATPase domain of all known remodeler structures contact nucleosomal DNA at SHL 2 (*2*). SMARCAD1, a single-subunit chromatin remodeler from the INO80 family is unique in that it sits astride SHL 0 and binds DNA at SHL - 1, while at the same time engaging extranucleosomal linker DNA. The ATPase domains of Iswi, which served as the template for the homology model of the SMARCAD1 ATPase domains, engages nucleosomal DNA at SHL -2 and at the neighboring gyre of the DNA superhelix through ATPase2 and 1, respectively, and is also anchored by interactions between ATPase2 and the H4 tail (fig. S8). Intriguingly the interactions between the H4 tail and ATPase2 are maintained in SMARCAD1, but the two domains appear to swivel around this anchor and translate by one SHL to interact with SHL -1, 0, and extranucleosomal DNA (Fig. S8). SMARCAD1 interactions with the nucleosome also differ from its closest ‘relative’ INO80 which contacts DNA at SHL 6 and SHL 3, highlighting the variability in interactions of the ATPase domains with their nucleosome substrates.

The ATP hydrolysis activity of SMARCAD1 is optimally triggered by the interaction with nucleosomes, and the energy is used to dissociate histones (presumably in their octameric form) from DNA. SMARCAD1 then deposits the histone octamer onto a new DNA segment. Because the two histone dimers are transferred at the same rate, and because SMARCAD1 brings H4 and H2B (and by extension, the histone dimers H3-H4 and H2A-H2B) into close proximity, we suggest that unlike other chromatin remodelers that exchange histone H2A-H2B dimers, SMARCAD1 is able to transfer the entire histone octamer. A similar activity has been suggested for the yeast homolog Fun30 (*23*) and for RSC (*5*). In the latter study, elegant kinetic analysis of histone octamer transfer suggests that RSC removes a histone octamer from DNA, and, in a second step, deposits it onto a new DNA segment. This model suggests that ATP-hydrolysis is only required to generate a ‘disrupted nucleosome intermediate’ (or, to remove histones from DNA), but not for the deposition of the histone octamer to form a new nucleosome, similar to the mechanism we propose here for SMARCAD1.

The two classes of particles described here are distinguished mainly by the amount of DNA bound on the SMARCAD1-distal side of the nucleosome. While in class 1 the DNA maintains all its contacts with the histone octamer, in class 2 DNA is peeled away from the nucleosome, making SHL +4 the first histone-DNA contact. The SMARCAD1 N-terminal domain, which is highly acidic and likely disordered, and therefore not defined in any of the current maps, projects towards this region. We speculate that this region may protect the DNA-binding surfaces of H2A-H2B that are exposed when the DNA is peeled away, similar in what was observed in the FACT-nucleosome complex (*34*).

Our functional and structural data suggest a mechanism (shown in fig. 5) in which SMARCAD1 engages nucleosome by interacting with extranucleosomal DNA and the DNA near SHL 0. In the presence of ATP, SMARCAD1 pushes DNA into the nucleosome and towards SHL 0. This destabilizes histone-DNA interactions, thereby allowing the N-terminal domain of SMARCAD1 to compete DNA away from SHL +4, and this step might be observed in class 2. As SMARCAD1 continues to hydrolyze ATP and peels off more DNA, the histones become more exposed, which eventually allows SMARCAD1 to completely remove them from DNA. Further work will focus on biochemically confirming this model, and higher resolution cryoEM analysis of this and other hypothetical intermediates will further describe SMARCAD1-nucleosome interactions.

**Fig. 5:**
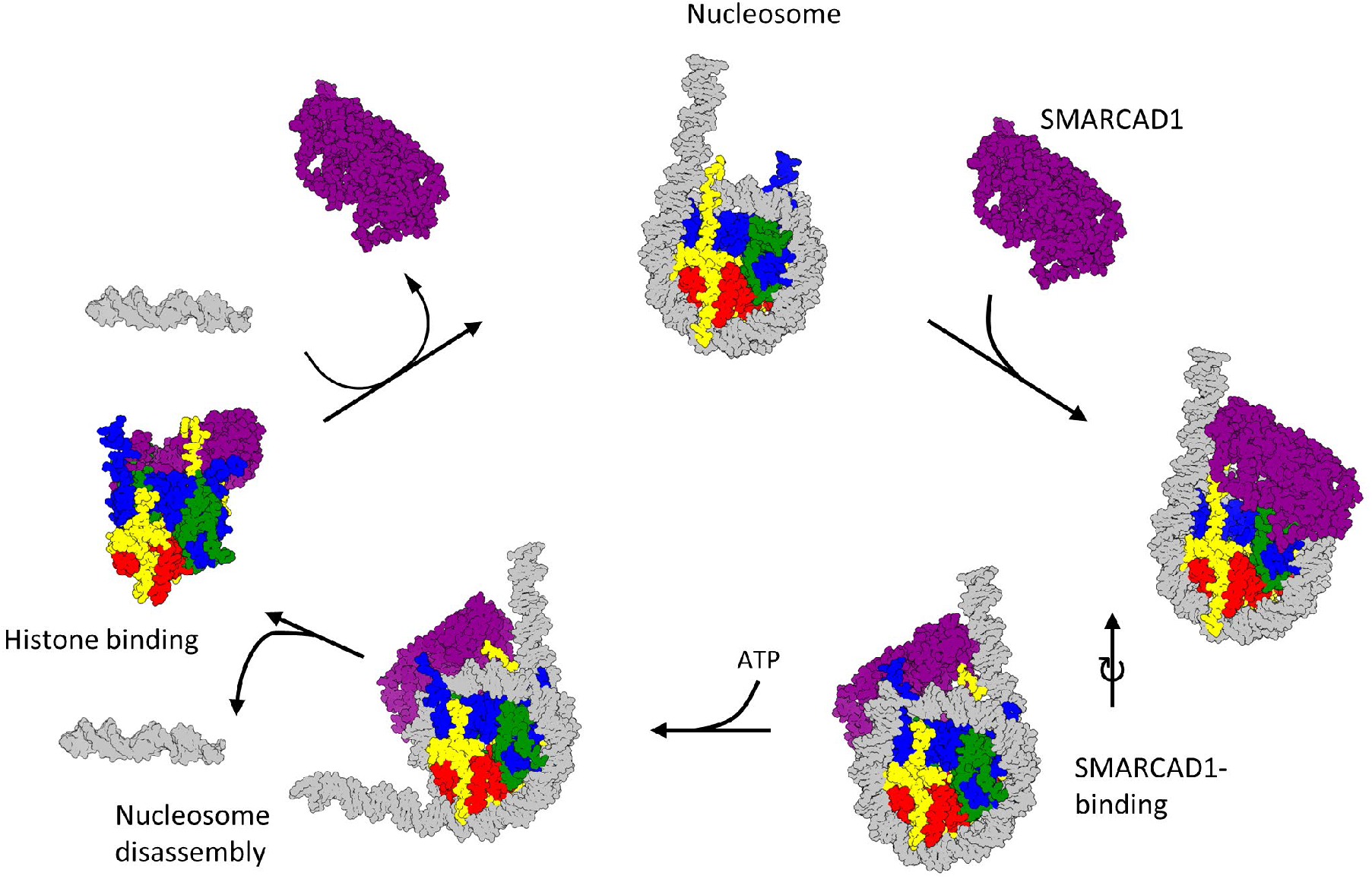
Proposed model for SMARCAD1 activity. SMARCAD1 first binds to the nucleosome on its proximal side by engaging extra-nucleosomal DNA and DNA at SHL 0 and SHL -1. SMARCAD1 then uses the energy of ATP-hydrolysis to peel DNA off the distal side of the nucleosome, allowing it to remove the histones. SMARCAD1 holds onto all four histones, possibly as a histone octamer, and deposits them back onto DNA to re-form a nucleosome.

Several remodelers interact with histone tails, most notably with the H4 N-terminal tail (*2*). Of particular interest, the H4 N-terminal tail relieves autoinhibition of Iswi, and this promotes ATP-hydrolysis (*35*). Here we show that SMARCAD1 interacts with both the H4 and H3 N-terminal tails. The H4 tail (but not the H3 tail) is required for ATP hydrolysis, while both are required for histone exchange onto a new DNA fragment. We speculate that SMARCAD1 can engage nucleosomal DNA (and free DNA) but is unable to hydrolyze ATP until it is properly situated on the nucleosome, as read out by proper interactions with the H4 tail. Further stabilization by the H3 tail then promotes histone eviction. The interactions with histone tails might thus provide a mechanism for regulating SMARCAD1 activity. Histone tails are extensively post-translationally modified, and this could further contribute to the regulatory mechanism.

## Acknowledgements

We thank the Janelia Research Campus Cryo-Electron Microscopy team, especially Zhiheng Yu, Shixin Yang, and Xiaowei Zhao for data collection. We also thank Garry Morgan at the University of Colorado Boulder Electron Microscopy core for help with grid screening.

## Author contributions

Conceptualization: JWM, KL

Methodology: JWM, KZ

Investigation: JWM, KZ

Visualization: JWM

Funding acquisition: KL

Project administration: KL

Supervision: KL

Writing – original draft: JWM, KL

Writing – review & editing: JWM, KZ, KL

## Data and materials availability

structures and maps will be deposited at the PDB and at EMPIAR. All data are available in the main text or the supplementary materials.

## Supplementary Materials

Figs. S1-S8

Table S1

Movie S1

## Materials and Methods

### Cloning and purification of SMARCAD1

A plasmid containing the human SMARCAD1 DNA sequence was purchased from DNASU. The sequence was PCR-amplified to add an N-terminal 6-His tag and restriction enzyme overhangs to allow cloning into the pACEBAC1 plasmid. Sf9 cells were infected with *Autographa californica* multiple nucleopolyhedrovirus (AcMNPV), and virus was amplified using the bac-to-bac system. Generally, expression was done in 300 mL Sf9 cells, and pelleted after 3 days expression. Cells were resuspended in lysis buffer (250 mM NaCl, 20 mM HEPES (pH 7.5), 2 mM TCEP, 1 mM AEBSF, 10% glycerol, 2 mM MgCl_2_) supplemented with a cOmplete Protease Inhibitor Cocktail (Roche Diagnostics) and 1500 units Benzonase (Millipore Sigma) per 300 mL Sf9 insect cells. Resuspended cells were lysed using a TissueLyser (Tekmar), incubated on ice for 10 minutes, sonicated, and spun at 31,000 RCF (Beckman JA-20 rotor) for 15 minutes, followed by purifications over a 5 mL nickel column and a 5 mL HiTrap-Q column. Pure fractions were mixed with 500 units Quick Calf Intestinal Alkaline Phosphatase (New England BioLabs) per 1 mg SMARCAD1 for 1 hour at room temperature. The sample was purified over a 1 mL nickel column followed by gel filtration over an S200 column. Samples were frozen in S200 buffer (100 mM KCl, 20 mM HEPES (pH 7.5), 2 mM TCEP, 1 mM AEBSF, 10% glycerol). Yield was in the range of 5-10 mg from 300 mL Sf9 cells.

### Histone refolding and nucleosome reconstitution

Human histones were purchased from the Histone Source at Colorado State University (Fort Collins) and refolded, and DNA was obtained as previously described (*36*). E63C H4 was labeled with a maleimide-Alexa488 fluorophore and T115C H2B was either labeled with a maleimide-Alexa488 or a maleimide-Atto647n fluorophore, as indicated (*37*). For tailless (TL) histone experiments, TLH3 (38-135) or TLH4 (20-102) were reconstituted the same as WT histones.

DNA for nucleosomes to be used for cryoEM were made from 10 mL PCR reactions using in-house purified Pfu-polymerase. The following DNA sequence was used (601 nucleosome positioning sequence in bold):

**ATCTGAGAATCCGGTGCCGAGGCCGCTCAATTGGTCGTAGACAGCTCTAGCAC CGCTTAAACGCACGTACGCGCTGTCCCCCGCGTTTTAACCGCCAAGGGGATTA CTCCCTAGTCTCCAGGCACGTGTCAGATATATACATCCGAT**ATCGGATCCTCTAG AGTCGACCTGCAGGCATGCAAGCTTGGCGTAATCATGGTCATAGCTGTTTCCTGTG.

Nucleosomes were reconstituted as described (*36*).

### Fluorescence Polarization

SMARCAD1 binding to nucleosomes was determined using a fluorescence polarization (FP) assay. Assays were done in binding buffer (20 mM Tris-Cl (pH 7.5), 2 mM DTT, 1 mM EDTA, 0.01% CHAPS, 0.01% NP40). Alexa-488 labeled H4 nucleosomes were used at a final concentration of 10 nM. SMARCAD1 was mixed with nucleosomes at increasing concentrations from 0-2500 nM in nucleosome binding buffer. The increase in FP-values as a function of SMARCAD1 concentration were measured in a BMG Labtech CLARIOstar plate reader. For the DNA-binding experiments, DNA was made by PCR amplifying 147 bp 601 sequence with primers containing Alexa-488 label (purchased from IDT). All FP binding curves were fit to the quadratic binding equation:

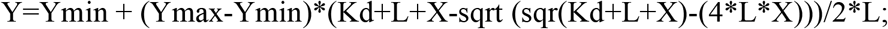

where Ymin and Ymax are the minimum and maximum FP Y signals respectively, Kd is the experimental binding constant, L is the ligand concentration (10 nM), and X is the SMARCAD1 concentration.

### ATP Hydrolysis Assay

We used an assay that couples ATP hydrolysis to NADH oxidation, using lactate dehydrogenase and pyruvate kinase (*4*). The reaction was done in ATP hydrolysis buffer (1 mM DTT, 4 mM MgCl2, 1 mM PEP, 12 µL LDH/PK enzyme mix (Sigma Aldrich), 50 mM HEPES (pH 7.5), 100 mM KCl, 0.7 mM NADH). 100 µL reactions were conducted by mixing 100 nM SMARCAD1, ATP Hydrolysis buffer, increasing amounts of activator, and then initiated by addition of 1 mM ATP. Reactions were measured in a BMG Labtech CLARIOstar plate reader, where NADH absorbance was measured at a wavelength of 340 nm. Rates were obtained from slopes of simple linear regression. Change in A340/s were converted to ATP-hydrolyzed ([M]/s) by using the NADH extinction coefficient (6330 M-1*cm-1). Background ATP-hydrolysis (SMARCAD1 without activator) was subtracted, then rates as a function of activator concentration were plotted and fit to the Michaelis-Menten equation:

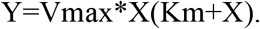

### Histone Exchange Assay

Tri-nucleosomes (30N60N60N30; the numbers refer to bp linker DNA, while N refers to 147 bp 601 nucleosome; Fig. 1C) were created in the same way as above, with H2B or H4 labeled with Atto647n. Histone exchange assays were done at 3 µM SMARCAD1, 62.5 nM tri-nucleosomes (equivalent to 225 nM individual nucleosomes), 1.5 µM acceptor 0N0 DNA, conducted in 2 mM DTT, 2mM MgCl_2_, 25 mM HEPES (pH 7.5), 50 mM KCl. Reactions were started with addition of 1 mM ATP, and at the appropriate timepoints were quenched with quench buffer (final concentration 68 mM EDTA and 16% glycerol). Samples were analyzed on 5% native TBE gels, imaged on Typhoon imager, then quantified with ImageQuant. Rates were calculated using a one-phase association equation:

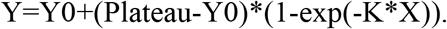

### FRET-based SMARCAD1-histone interactions

To determine if SMARCAD1 binds to histones H2A-H2B and H3-H4 simultaneously in the absence of DNA, we used a FRET-based approach that allows us to monitor (H3-H4) interactions with (H2A-H2B). Refolded Alexa488-H3-H4 and Atto647n-H2A-H2B histones were mixed at low concentration (10 nM) in 2 mM DTT, 2 mM MgCl2, 50 mM HEPES (pH 7.5), 0.01% CHAPS, 0.01% NP40. SMARCAD1 was added at increasing concentration (0-125 nM). The increase in FRET-values as a function of SMARCAD1 concentration was measured in a BMG Labtech CLARIOstar plate reader, where highest SMARCAD1 titration points were arbitrarily set at 50% fluorescent intensity. Binding constants were calculated using a hyperbolic binding equation:

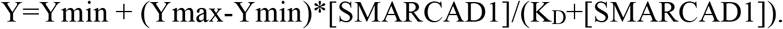

### SMARCAD1 *de novo* nucleosome assembly assay

To monitor if SMARCAD1 assembles nucleosomes *de novo*, we used an assay that monitors incorporation of histones onto DNA as more SMARCAD1 is added. SMARCAD1 (indicated concentrations) was mixed with H3-H4 (50 nM) and H2A-H2B (Atto647n-H2B; 50 nM) for 15 minutes in 2 mM DTT, 100 mM KCl, 50 mM HEPES (pH 7.5), 10% glycerol, 2 mM MgCl2. 147 bp DNA (50 nM) was added (with 1 mM ATP when indicated) for 15 minutes. Samples were quenched with 16% glycerol, 68 mM EDTA, and 2.7 µg pUC19 plasmid. Reactions were analyzed by a 5% native TBE gel, imaged on a Typhoon imager, and then quantified with ImageQuant.

### FP-based quantification of SMARCAD1-histone interactions

We used the FP assay to determine if SMARCAD1 binds H3-H4 and H2A-H2B individually in the absence of DNA. Alexa488-H3-H4 (5 nM) or Alexa488-H2A-H2B (5 nM) in 2 mM TCEP, 2% glycerol, 50 mM HEPES (pH 7.5), 0.01% CHAPS, 0.01% NP40, and 100 mM KCl was combined with increasing concentrations of SMARCAD1 (0-2.5 µM). The increase in FP signal as a function of SMARCAD1 concentration was determined in a BMG Labtech CLARIOstar plate reader, where no SMARCAD1 H3-H4 signal was arbitrarily set at an FP value of 100 mP. Binding constants were calculated using a hyperbolic binding equation:

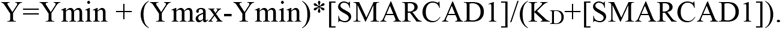

### CryoEM SMARCAD1-nucleosome sample preparation

To prepare SMARCAD1-nucleosome complex for cryoEM, we first reconstituted a 0N70 nucleosome using published procedures (*36*). We then combined SMARCAD1 (final concentration 16 µM) with 0N70 nucleosome (2 µM) in S200 buffer. For crosslinking, GraFix buffers with top buffer (10% glycerol, 50 mM NaCl, 10 mM HEPES; pH 7, 1 mM MgCl2, 500 µM ADP, 8 mM NaF, and 1 mM BeSO4) and bottom buffer (30% glycerol, 50 mM NaCl, 10 mM HEPES (pH 7), 1 mM MgCl_2_, 500 µM ADP, 8 mM NaF, 1 mM BeSO4, and 0.15% glutaraldehyde) were used. A 10-30% glycerol gradient was generated, and 200 µL SMARCAD1-nucleosome sample was added to the top. Samples were spun at 28K RPM in an sw40-TI rotor (Beckman) for 20 hours in an ultra-centrifuge. Fractions were collected, analyzed by TBE and SDS PAGE, and fractions containing complex were collected. They were then dialyzed into 50 mM NaCl, 10 mM Tris (pH 8), 2 mM MgCl_2_, 1 mM ADP, 16 mM NaF, 2 mM BeSO_4_, and 3 mM DTT (*24, 25*).

### CryoEM grid preparation and data collection

4 µL of dialyzed sample was deposited onto glow-discharged C-flat Au 1.2/1.3 grids (Electron Microscopy Sciences), and flash frozen using a manual plunger. Grids were screened at the University of Colorado Boulder CryoEM facility using a Tecnai T20 microscope with a K3 electron detector. High quality grids were then sent to the Janelia Research Campus cryoEM facility for data acquisition. The complex was imaged at a magnification of 64000x on an FEI Titan Krios (300 kV), equipped with a Gatan K3 summit direct detector. Pixel size was 1.065 Å. The movies were captured in super resolution mode with an electron dose rate at 7.8 electrons per pixel per second for 5.593s at 0.13 s/frame. The defocus range was -1.2 to -2.5 µm. A total of 11,160 images were collected.

### CryoEM processing and model building

The data was processed in two separate steps. In the first round of processing, 2,408 movies were patch motion corrected and patch CTF corrected in Cryosparc version 2.4 (*38*). Particles were then picked with the Cryosparc blob picker and extracted at a box size of 256 A^2. Extracted particles were subjected to 2D classification, then good classes were used as templates in the Cryosparc template picker. The template picker resulted in 1,513,927 particles. These were subjected to 2D classification in Cryosparc, and bad particles were removed, resulting in 440,777 particles. These particles were then transferred to Relion version 3.0 for further processing (*39*). Using an initial model from 10,184 particles, all particles were subjected to two rounds of 3D classification. After 3D classification, we had 320,523 particles present in one class. These particles could not be cleaned up further even when initial the model was low pass filtered to 30 Å. To increase the resolution of the SMARCAD1 density, a mask was created that only encompassed the SMARCAD1 region. With this mask, particles were further subjected to two rounds of 3D classification without image alignment. Best particles were then subjected to 3D refinement, which resulted in a final resolution of 6.49 Å (FSC = 0.143). FSC curves were determined with the 3D FSC server (*40*).

In the second round of processing, 5,395 movies were patch motion corrected and patch CTF corrected in Cryosparc. Using the same templates from the first round of processing, a total of 3,282,892 particles were picked. These were subjected to one round of 2D classification in Cryosparc, which resulted in 943,877 particles. These were brought to Relion, where they underwent two rounds of 3D classification. A third round of 3D classification was performed with local angular searches, which revealed a new SMARCAD1-nucleosome class, showing DNA peeled away from the histone core. This class was auto refined to a resolution of 4.4 Å (FSC = 0.143).

For the nucleosome, model building was performed by placing the human nucleosome (PDB 2CV5) into the density. We manually built the DNA into the density using Coot (*26*). As noted in the results, the density for SMARCAD1 is at lower resolution than the nucleosome due to its high flexibility. Therefore, it was difficult to manually build the SMARCAD1 structure. To aid in SMARCAD1 model building, we first used Swiss-Model to predict a structure of SMARCAD1 using Iswi (PDB 5JXR) as a template (*27*). The predicted structure was then docked into the EM map in chimera (*28*). To better fit the model into the density map, the structure and map were run in Flex-EM (*29*). The model was then simulated in MDFF (using the NAMD engine and visualized in VMD) and then further processed in ISOLDE to further increase model to map fitting (*30-33*).

**Fig. S1.**
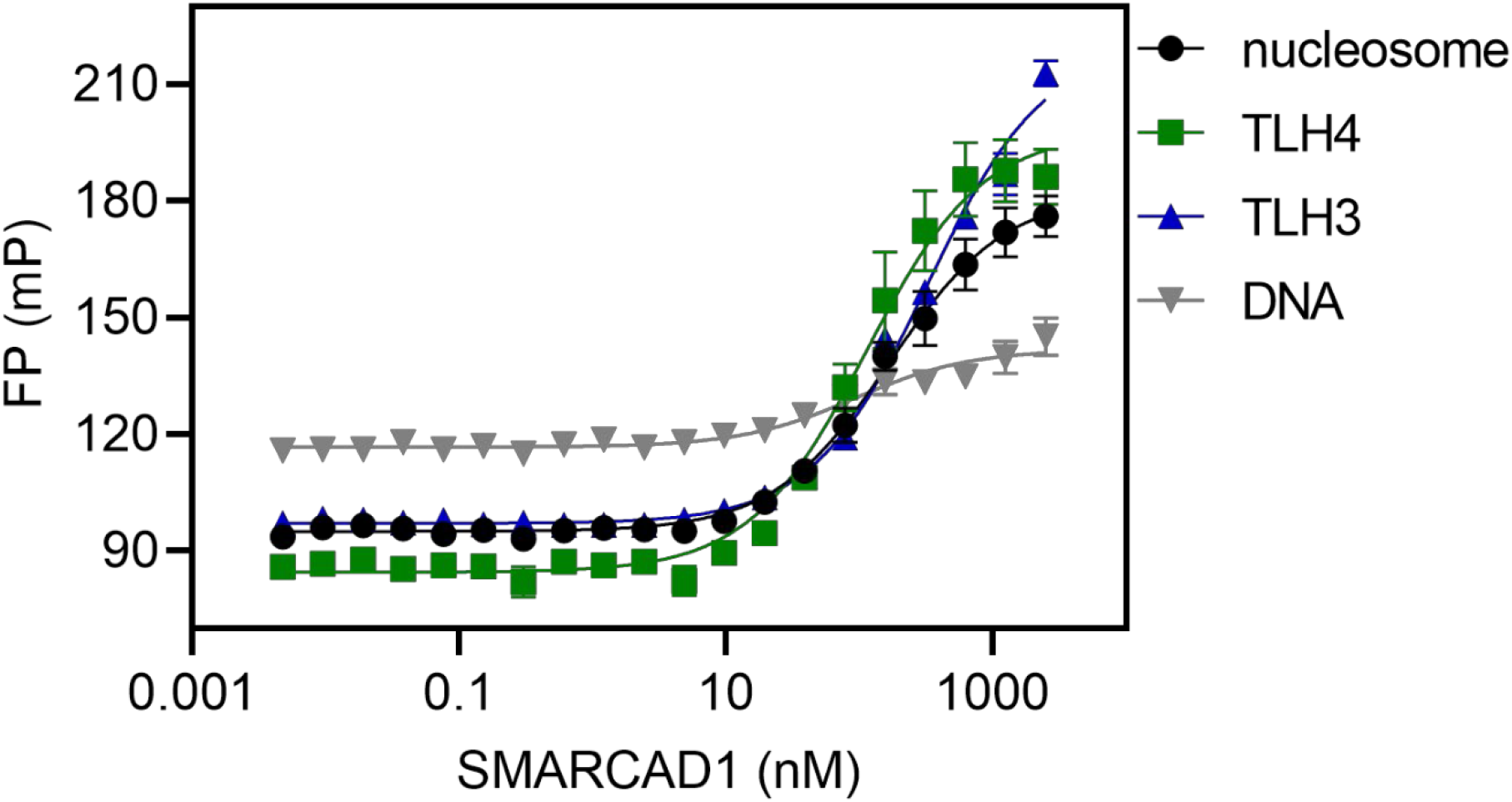
Fluorescence polarization raw data (figures 1A and 4C). The same data as shown in fig. 1A and 4C, without normalization. Differences in FP (mP) signal varies in magnitude when using nucleosomes or free DNA, due to different size and shape of the substrates.

**Fig. S2.**
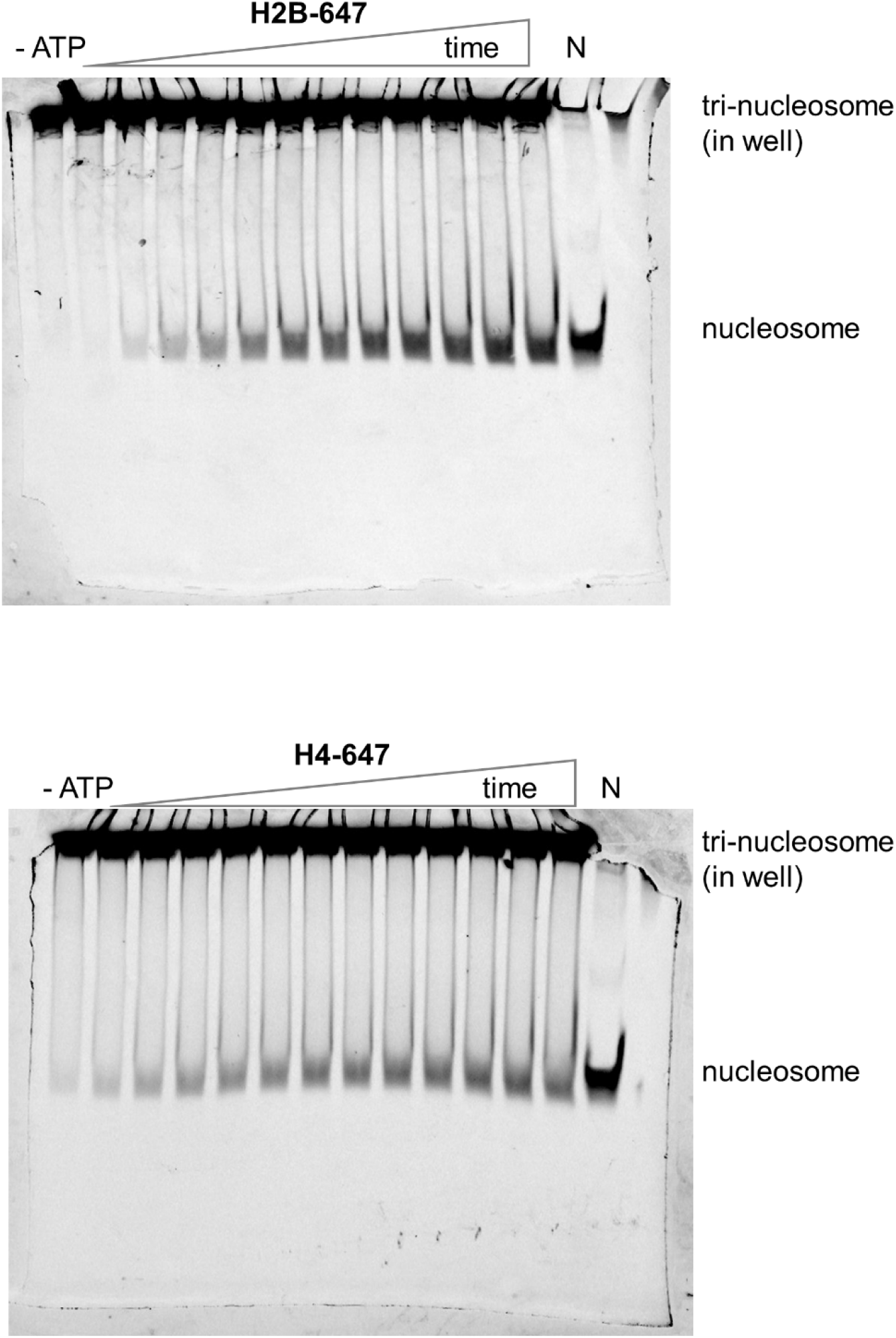
Uncropped raw gel files for the histone exchange assay shown in figure 1D. 5 % native PAGE in 0.2x TBE. Time points were 0, 60, 120, 180, 240, 300, 420, 540, 600, 720, 900, and 1,800 seconds.

**Fig. S3.**
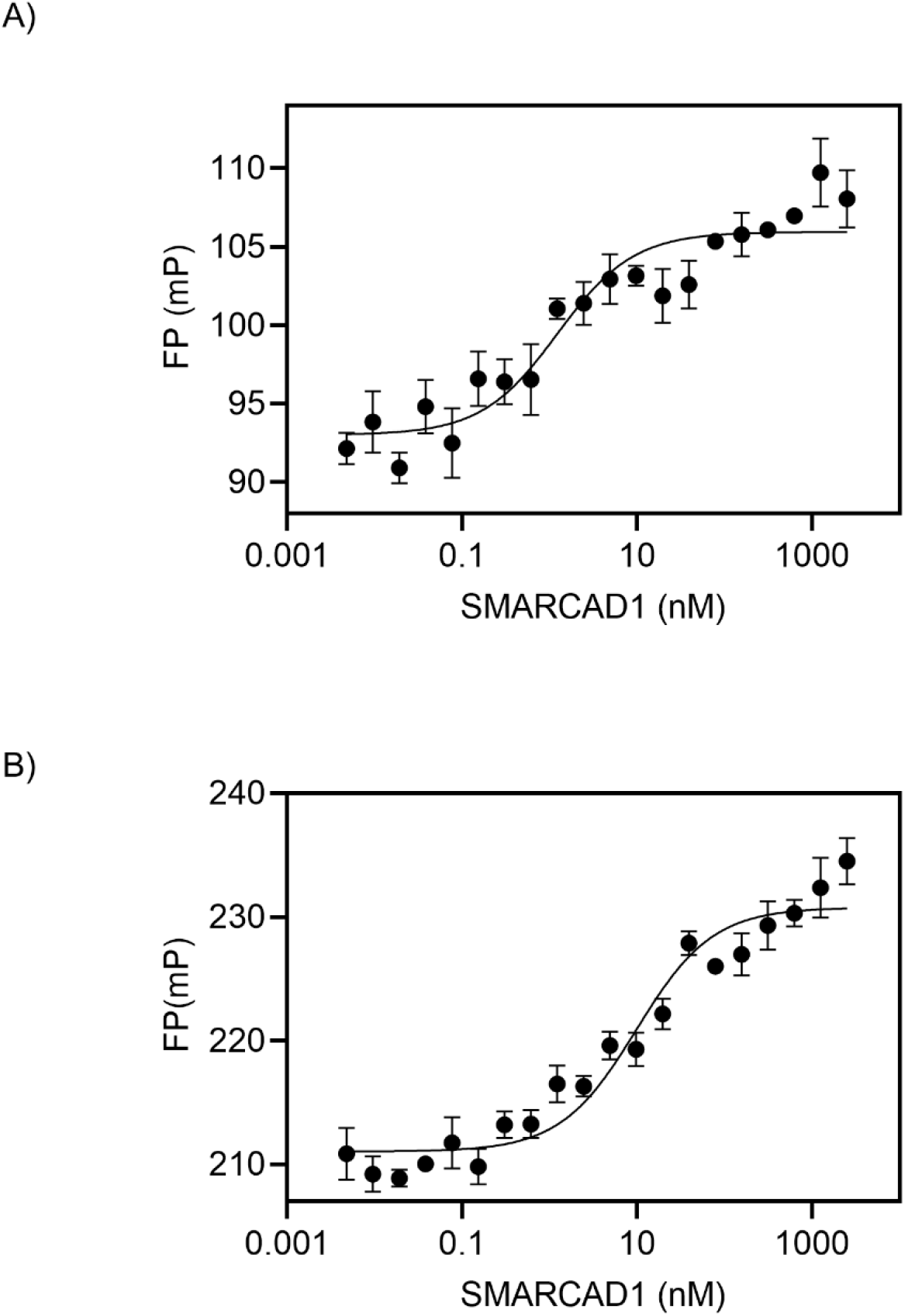
SMARCAD1 binds histones H3-H4 and H2A-H2B independently (Fluorescence Polarization). **A)** SMARCAD1 (0-2,500 nM) binds H3-H4 (Alexa-488 H4, 5 nM) in absence of H2A-H2B or DNA with high affinity (K_D,H3H4_ = 1.2 ± 0.4 nM). K_D_ and s.e. from 4 replicates. **B)** SMARCAD1 (0-2,500 nM) binds H2A-H2B (Alexa-488 H2B, 5 nM) in absence of H3-H4 or DNA with high affinity (K_D,H3H4_ = 10 ± 2 nM). K_D_ and s.e. derived from 4 replicates.

**Fig. S4.**
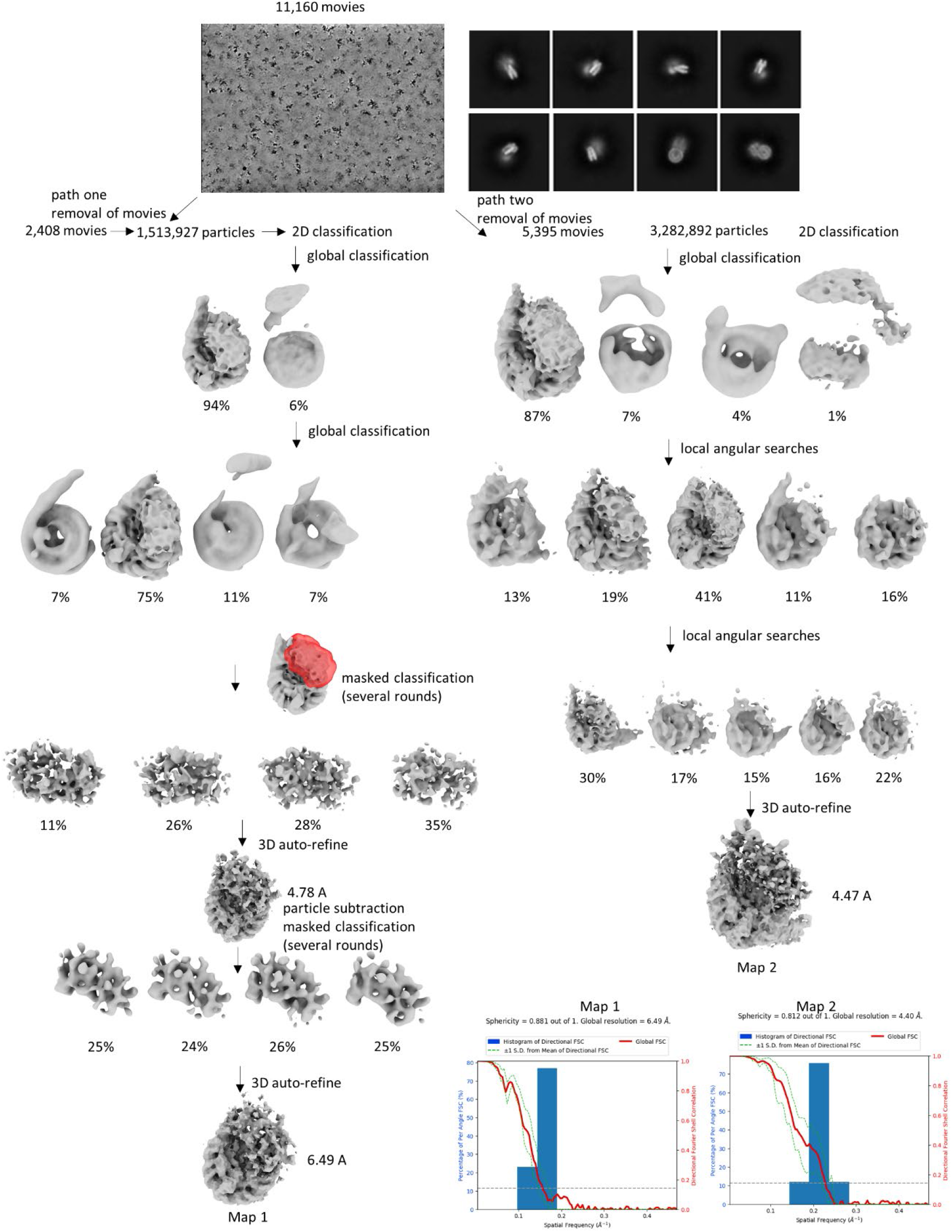
CryoEM analysis workflow. FSC curves of map 1 and map 2 determined with 3D FSC server, overall FSC curves and resolutions written on graph (FSC = 0.143)

**Fig. S5.**
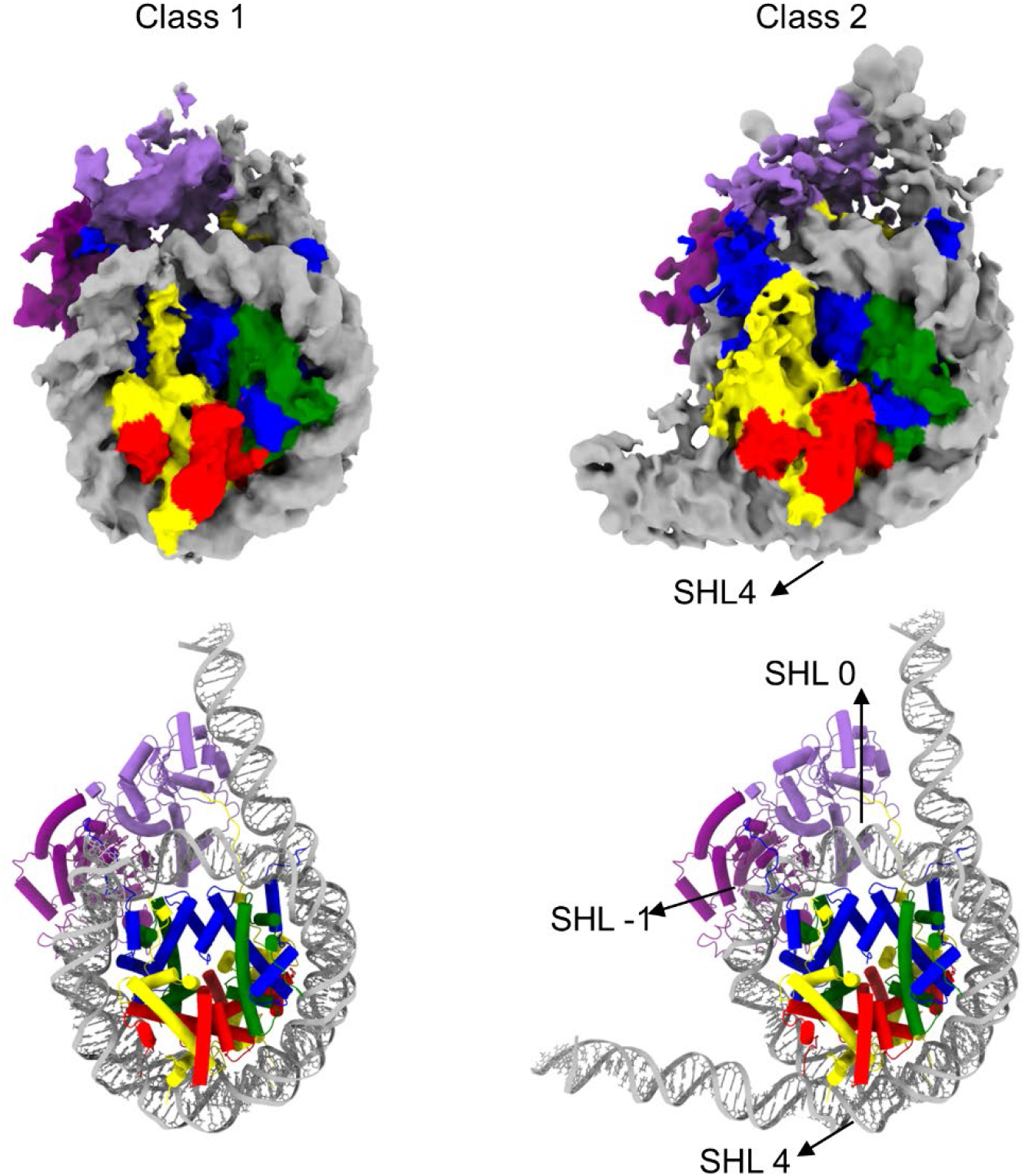
Comparison of map 1 and map 2 reveals that SMARCAD1 peels DNA off the distal side starting at SHL 4, exposing the histones, which ultimately may result in nucleosome disassembly. Color code as in fig. 3B. To improve clarity, the ‘surface dust’ command was used in ChimeraX to remove noise from maps.

**Fig. S6.**
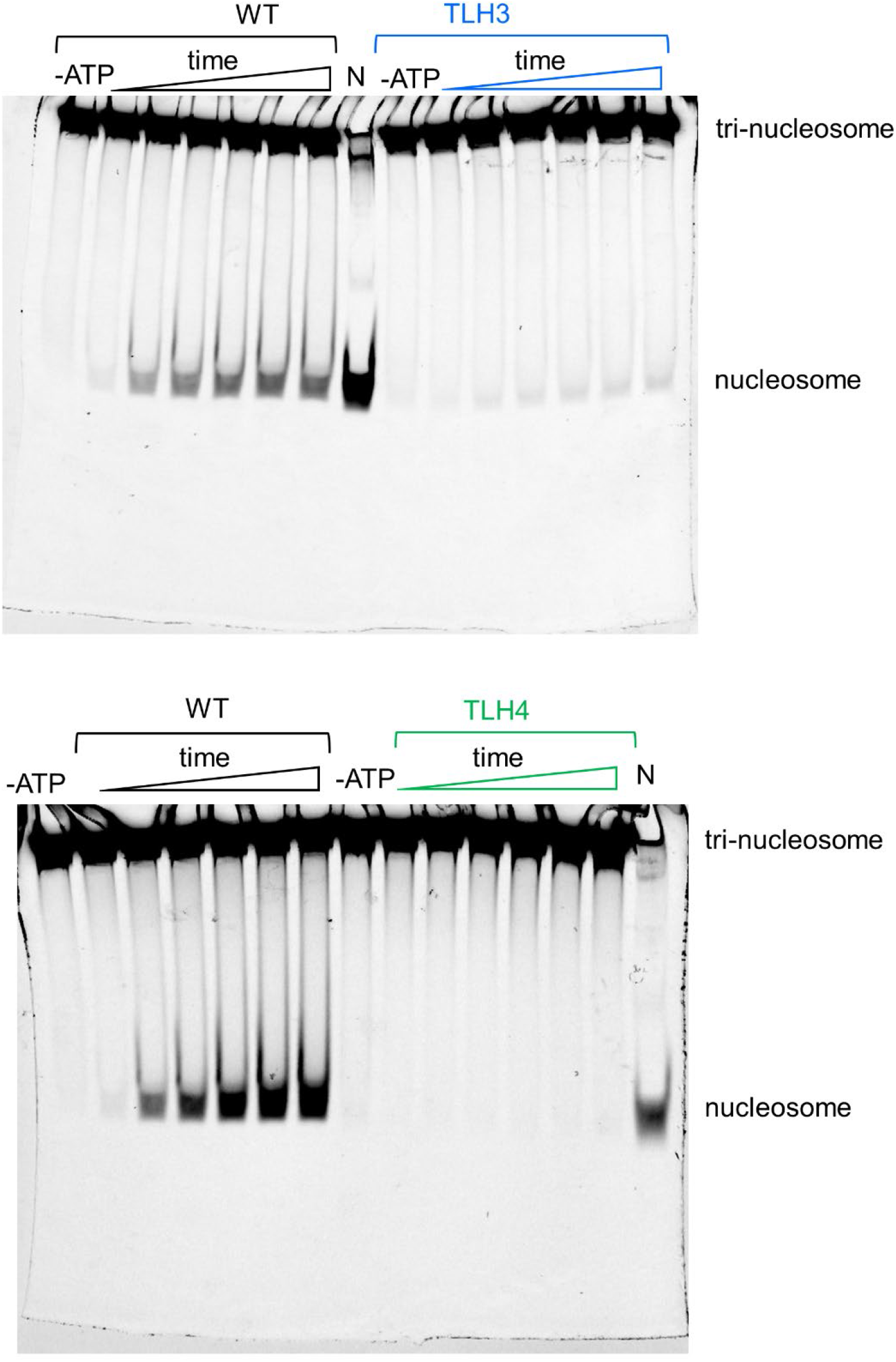
Uncropped gels from figure 4E. Time points were 0, 60, 18, 300, 540, 900, 1800 seconds.

**Fig. S7:**
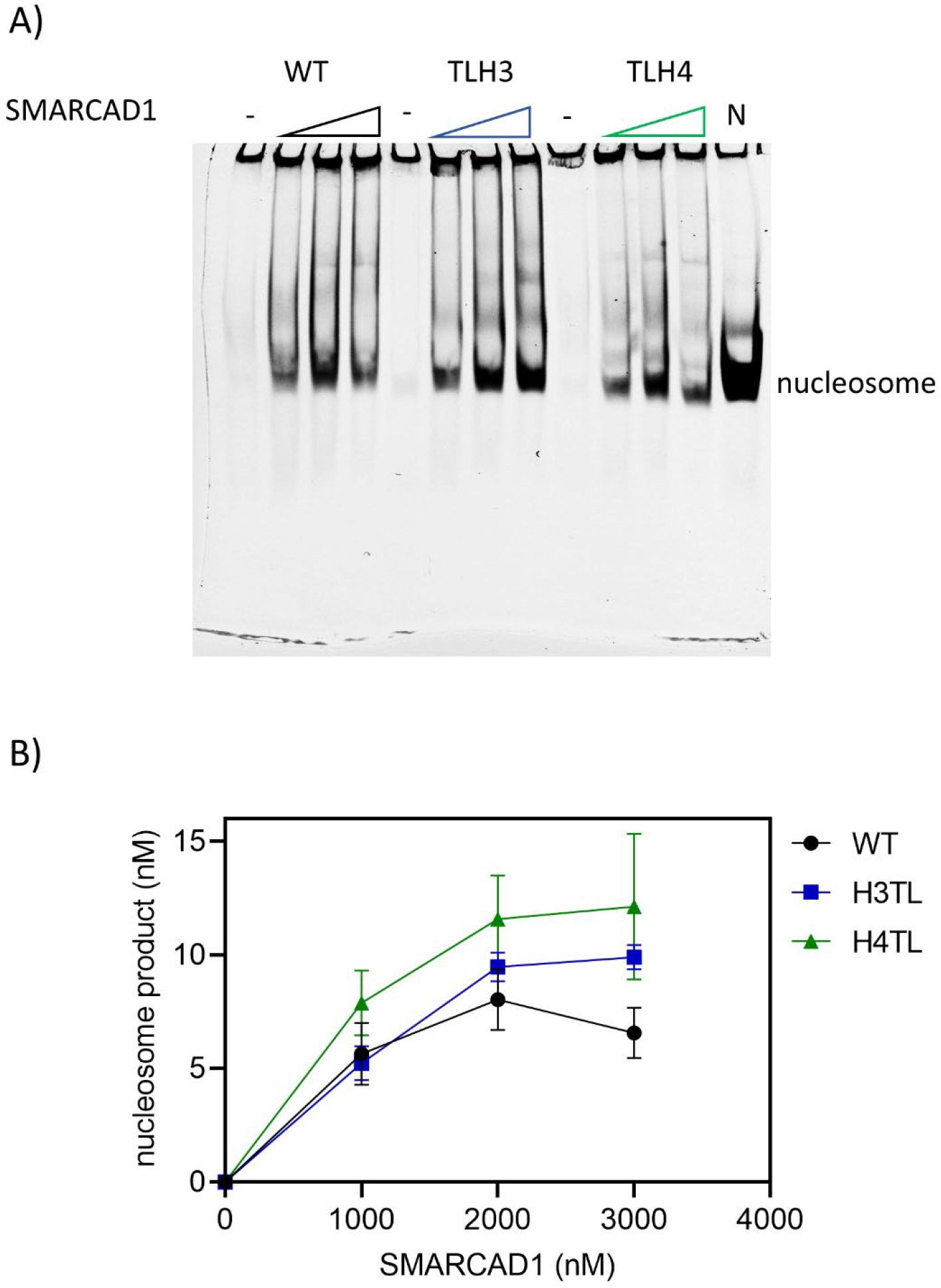
Histone tails are not required for nucleosome assembly from histones. Using the assay described in figure 2C, H3-H4 and H2A-H2B (50 nM; 647-H2B) were mixed with SMARCAD1 (0-3 μM) for 15 minutes followed by addition of 147 bp DNA (50 nM) and a 15 minute incubation. Reactions were quenched with pUC-19 plasmid DNA and then analyzed on a 5% native TBE gel. In the absence of SMARCAD1, little nucleosome assembly occurs; however, after addition of SMARCAD1, there is an increase in nucleosome assembly for WT, TLH3, and TLH4 histones. **B)** Quantification of gel from a) confirms that SMARCAD1 assembles WT, TLH3, and TLH4 histones into nucleosomes. s.e. bars in graph are from four replicates.

**Fig. S8:**
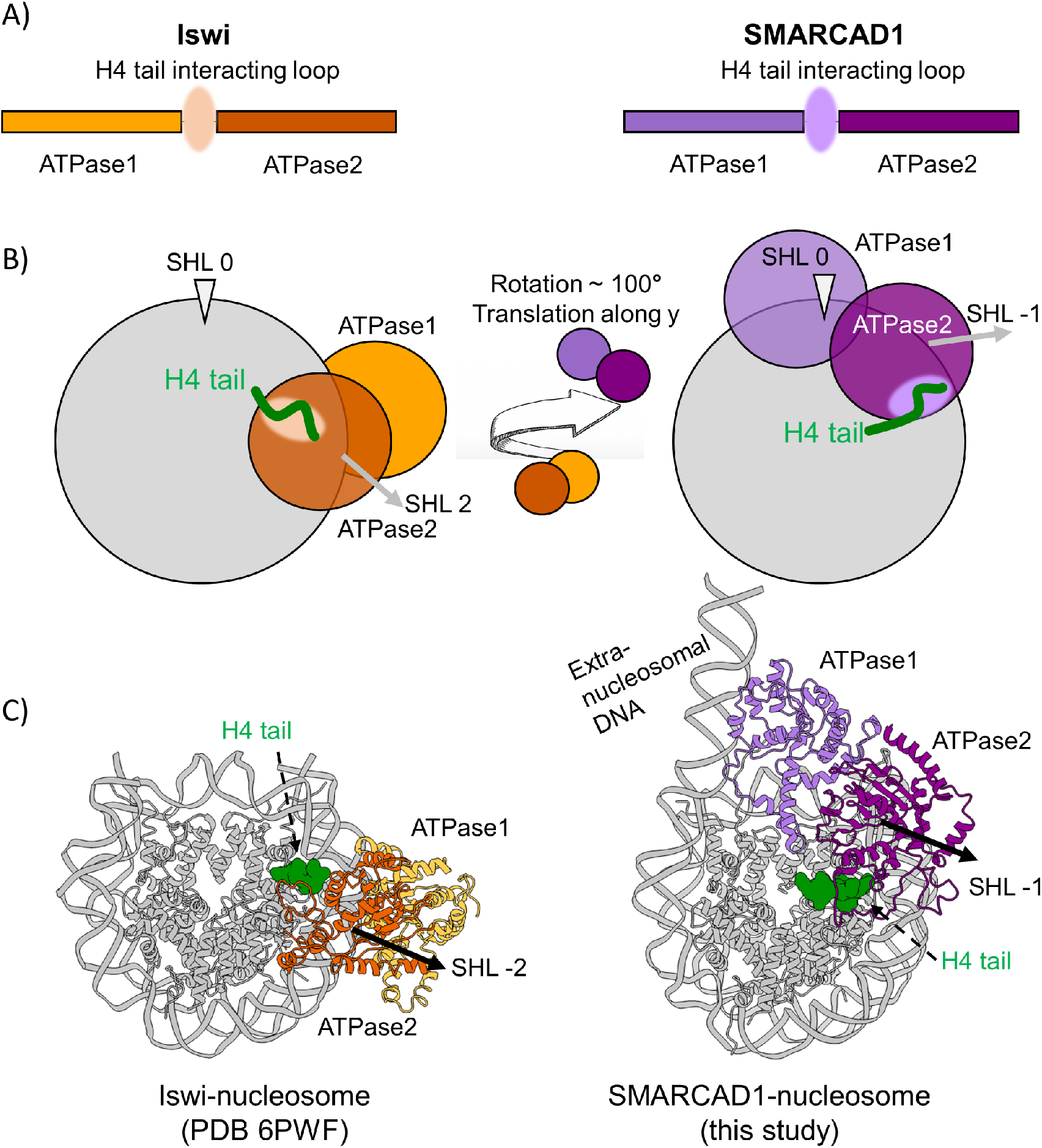
Comparison of nucleosome interactions between the catalytic domain of Iswi (PDB 6PWF) and SMARCAD1 (this study). **A)** Both Iswi and SMARCAD1 have similar catalytic domain architecture. Two ATPase domains are separated by a linker (shaded oval) that interacts with the H4 N-terminal tail. **B)** Cartoon representation. Iswi ATPase2 binds SHL -2 and the H4 tail, while ATPase 1 is behind ATPase2 where it interacts with the neighboring gyre of DNA at SHL +6. The ATPase domains of SMARCAD1 pivot around the H4 anchor point by ∼100 degrees relative to Iswi, where the ATPase2 domain binds to SHL -1, putting ATPase1 in position to bind extranucleosomal DNA. **C)** Structures indicating histone tail and DNA interactions of the two ATPase domains of the two remodelers.

**Table S1.** Data collection and refinement parameters

|  | **Class 1**<br>**(PDB xxxx)** | **Class 2**<br>**(PDB xxxx)** |
| --- | --- | --- |
| **Data collection and processing** |  |  |
| Magnification | 64,000 | 64,000 |
| Voltage (kV) | 300 | 300 |
| Electron exposure (e-/Å<sup>2</sup>) | 60 | 60 |
| Defocus range (μm) | 1.2-2.5 | 1.2-2.5 |
| Pixel size (Å) | 1.065 | 1.065 |
| Symmetry imposed | C1 | C1 |
| Initial particle images (no.) | 1,513,927 | 3,282,892 |
| Final particle images (no.) | 20,135 | 130,885 |
| Map resolution (Å) | 6.49 | 4.40 |
| FSC threshold | 0.143 | 0.143 |
| Map resolution range (Å) | 4.6-11.6 | 4.2-10.4 |
| **Refinement** |  |  |
| **Initial models used (PDB code)** | 2CV5, 5JXR | 2CV5, 5JXR |
| **Model composition** |  |  |
| Non-hydrogen atoms | 17479 | 17479 |
| Protein residues | 1312 | 1312 |
| Nucleotides | 340 | 340 |
| **R.m.s. deviations** |  |  |
| Bond lengths (Å) | 0.004 | 0.011 |
| Bond angles (°) | 0.681 | 0.725 |
| **Validation** |  |  |
| MolProbity score | 2.40 | 2.27 |
| Clashscore | 22.42 | 19.11 |
| Poor rotamers (%) | 0.00 | 0.00 |
| **Ramachandran plot** |  |  |
| Favored (%) | 89.64 | 91.89 |
| Allowed (%) | 10.28 | 7.96 |
| Disallowed (%) | 0.08 | 0.15 |

**Movie S1**. cryoEM maps and models of SMARCAD1-nucleosome complex. Specific regions are focused upon to summarize important interactions for the two different models.

